# Identification and characterization of *Porphyromonas gingivalis* TonB

**DOI:** 10.1101/2025.11.18.689179

**Authors:** Soheil Rostami, B. Ross Belvin, Janina P. Lewis

**Author notes:** For Correspondence: Janina P. Lewis, PhD, Virginia Commonwealth University, Philips Institute for Oral Health Research, 521 North 11^th^ Street, Room 401, Richmond, VA 23298, USA. These authors contributed equally to the manuscript (Soheil Rostami, B. Ross Belvin). Data are available via ProteomeXchange with identifier PXD033524.

## Abstract

*Porphyromonas gingivalis* is one of the major bacterial pathogens responsible for the initiation and progression of periodontal disease. The bacterium codes for multiple TonB-dependent receptors required for acquisition of nutrients such as heme and vitamin B12, although the identity of a TonB energy transducer has yet to be identified. Here we identify a potential TonB protein encoded by PG0785, generate a deficient strain, and characterize its biological significance. Bioinformatics analysis reveals that the PG0785 has unique features confined to the Cytophaga-Flavobacterium-Bacteroides (CFB) group of bacteria but shares similarities in the C-terminal domain (CTD) to well characterized proteins from *Helicobacter pylori* and *Escherichia coli.* Similarity at a protein level, as well as the genomic locus organization, was identified with the TonB proteins characterized in other *Bacteroidota* species. Loss of PG0785 led to significant alteration of the proteome: proteins mediating gene regulation, protein translation, protection against reactive radical species (oxygen and nitrogen), and glycosylation were up-regulated and polysaccharide export, efflux protein, and lipoproteins were downregulated. Furthermore, phenotypic studies showed that a mutant lacking PG0785 cannot accumulate heme on its surface and is deficient in gingipain protease activity. In addition, reduction of the capsular layer was detected in the TonB-deficient strain. Thus, while the mutant interacted and invaded eukaryotic cells at much higher levels than the wild type, it had significantly reduced ability to survive with host cells.

**Importance:** The TonB system is indispensable for energy transduction in Gram-negative bacteria and is mainly associated with nutrient uptake. Although well investigated in Proteobacteria, the knowledge regarding the system in the CFB group of bacteria is lagging. In contrast to Proteobacteria, the intestinal Bacteroidota have multiple TonBs and their novel involvement in sugar utilization has been demonstrated. This study shows that the oral Bacteroidota, *Porphyromonas gingivalis*, also has TonB but unexpectedly, it is not involved in iron homeostasis or nutrient acquisition but rather in processes involved in maturation of proteases and cell surface remodeling. As maintenance of outer membrane integrity is mainly associated with Tol motor our study shows possible crosstalk between the energy transducers. Also, this study further demonstrates the functional versatility of the TonB system depending on the environmental niche and the metabolic requirements of the bacteria.

## Introduction

*Porphyromonas gingivalis* is a Gram-negative, rod-shaped, anaerobic bacterium that belongs to the Cytophaga-Flavobacterium-Bacteroides (CFB) group of bacteria. Biological studies have shown that *P. gingivalis* is one of the main etiological agents involved in initiation and progression of periodontal disease. It can be present in the oral cavity both extracellularly as well as internalized by host cells.

As for a Gram-negative bacterium, an efficient outer membrane transport system is indispensable for *P. gingivalis’* fitness. While diffusion through the outer membrane is one of the mechanisms for transport, *P. gingivalis* relies on larger molecules for its growth that include peptides^1^ and heme^2 3 4^. Heme is an essential nutrient for the bacterium and extracellular *P. gingivalis* acquires it primarily from the host’s heme containing proteins^4 5 6 7^. *P. gingivalis’s* proteases were shown to play a role in heme acquisition through degradation of heme-containing proteins such as hemoglobin^7^.

The TonB/ExbB/ExbD complex plays a major role in the passage of large and complex nutrients through the double-membrane of Gram-negative bacteria. The inner membrane of the bacterium generates a proton motive force (pmf) that is transferred via ExbB/ExbD to TonB^8^. TonB spans the periplasm and interacts with outer-membrane receptors. This interaction leads to the removal of C-terminal plug domain from inside of a membrane channel, which then allows for nutrients to be taken up through the receptor. The genome of *P. gingivalis*, and closely related bacteria *Tannerella spp.* and *Prevotella spp.*, code for a significant number of TonB-dependent transporters. Peptides are proposed to be transported via the novel RagAB transport system and heme transported through the Hmu system^1 9 10^. However, the genomes of these bacteria code for a significant number of TonB-dependent transporters with yet unknown or hypothetical transport function. These systems are dependent on and require TonB to provide the energy for transport. Although it is evident that TonB would be critical for growth of the organism, currently no *P. gingivalis* TonB proteins have been experimentally identified and characterized.

The genome of *P. gingivalis* W83 codes for a putative TonB protein (PG0785), however, the gene was not identified as iron regulated in our previous studies^3^ pointing to its unusual role in this bacterium. To gain insight into the function of the putative *P. gingivalis* TonB, we generated a mutant that was deficient in PG0785 and characterized the mutant in terms of its contribution to growth, metal content, protease activity, cell surface composition and survival with host cells. Also, we determined the whole cell proteome to gain an insight into the extent of the modifications present in the mutant cells when compared to the wild type.

## Materials and Methods

### Bacterial growth conditions

*Porphyromonas gingivalis* strain W83 was grown anaerobically at an atmosphere of 10% H_2_, 10% CO_2_ and 80% N_2_ at 37° C. Bacterial liquid cultures were prepared in brain-heart infusion broth (BHI; Difco Laboratories, Detroit, MI) with hemin (5 µg/ml; Sigma, St. Louis, MO), yeast extract (5 mg/ml), cysteine (1 mg/ml; Sigma, St. Louis, MO) and vitamin K3 (1 µg/mL; Sigma, St. Louis, MO).

### Mutagenesis of PG0785 and complementation

*P. gingivalis* W83 cells were grown anaerobically on blood agar plates (TSA II, 5% sheep blood; BBL, Cockeysville, MD). From freshly inoculated blood plates an overnight culture was prepared by inoculating 3 ml of BHI broth. Following overnight (ON) growth it was diluted 1:10 in broth and cultures were grown until an optical density (OD) of approximately 0.3 was attained. Bacterial cells were then pelleted and washed twice with electroporation butter (10% Glycerol; Fisher BioReagants). The cells were then used for generation of mutant strain. 10 µL of a suicide vector composed of pCR2.1 vector with the PG0785 interrupted at the *Nru*I site (located at the 82bp of the gene) with an *ermF-ermAM* cassette^11^ was mixed with *P. gingivalis* W83 electroporation cells, transferred into electroporation cuvettes (BIO-RAD, Hercules, CA) and electroporated at 2,500 mV using a Gene-Pulser (BIO-RAD, Hercules, CA). The cuvette was then transferred to an anaerobic chamber and 500 µL of BHI was added to the cuvette. Following overnight incubation, the culture was then spread on blood agar plates containing clindamycin (0.5 µg/ml) and incubated for 7 days. Colonies appearing on the plates were then screened using PCR and gel electrophoresis was used to assess the size of the PCR fragment. The mutant strain was designated V3128. For complementation, the PG0785 gene and the upstream region corresponding to its promoter was cloned into the pG108 vector using Gibson assembly^12^. The subsequent pG108K-*tonB* construct was electroporated into the V3128 mutant strain using the same settings and protocol as above. The overnight culture was plated onto blood agar plates containing tetracycline (0.5 µg/ml) and incubated for 5-6 days. Tet resistant colonies were re-plated on double antibiotic plates (Clin/Tet) and the presence of the plasmid was additionally checked using a plasmid miniprep kit. The complemented strain was denoted as V3128C.

### Protease assay

W83, V3128 and V3128C strains were grown overnight in BHI broth. The bacteria were harvested and pellets were suspended in PBS. Protease activity of both bacterial cells and culture supernatants was examined

#### Arginine-Specific Protease Assay

Buffer consisting of Tris-HCl (50mM, pH 7.4), L-Cysteine (5mM), CaCl_2_ (5mM), DTT (5mM), and Nα-Benzoyl-DL-arginine 4-nitroanilide hydrochloride (BAPNA) (1mM) was mixed with bacterial samples. Reactions were performed in a 96 well plate in a volume of 200 µL. To initiate the reaction, 5.0x10^6^ bacterial cells in PBS or 50 µL of culture supernatant was added to reaction wells. Reaction rates were monitored using an H1 BioTek plate reader measuring absorbance (Abs) at 405 at 37°C under aerobic conditions over a period of 180 min.

#### Lysine-Specific Protease Assay

Buffers consisting of Tris-HCl (50mM, pH7.4), L-Cysteine (5mM), 5mM DTT, and 5mM CaCl_2_, TGPLpNA (N-p-Tosyl-Gly-Pro-Lys-pNA) (0.5mM) were mixed with bacterial cells in a 96 well plate in a volume of 100 µL. 5.0x10^6^ bacterial cells in PBS or 25 µL of cell culture supernatant were added to reaction wells to initiate the reaction. The Lys-X protease activity was determined as above.

### Interaction of P. gingivalis with eukaryotic cells

The study of *P. gingivalis* – host interaction was done as described previously^13^. Normal oral keratinocytes (NOKs) and HL60 cells were used in this study. NOKs were cultured in keratinocyte media (ScienCell). HL60 cells were cultured in RPMI supplemented with 20% FBS. All cells were grown in a 5% CO_2_ incubator.

*P. gingivalis* W83, TonB-deficient *P. gingivalis (*V3128), and mutant complement strain (V3128C) were suspended from cultures grown on blood plates in 4 ml of BHI broth to an OD_660_ of 0.5. Bacteria were pelleted by centrifugation at 8000 rpm for 10 minutes, washed, and suspended in appropriate cell culture media.

NOKs were passaged and plated in 6 or 12 well plates. The next day, plates were moved into the anaerobic chamber, washed, and bacteria were then added to each well with a multiplicity of infection (MOI) of 100:1 (bacteria: eukaryotic cells). The infected cells were incubated for 30 minutes to 1 hour and then washed 3 times with PBS. Immediately, 2 mls of BHI broth with 1% saponin (Riedel-de Haen, Germany) was added to each well and the mixture was incubated for 15 minutes. To determine the number of adhered and internalized bacteria, the host cell lysis mixture from each well was serially diluted and plated on blood agar plates.

To test for internalization, after infection and incubation, cells were washed and a combination of metronidazole (300 µg/mL) and gentamicin (400 µg/ml) was added to the cells for 1 hour to kill any internalized bacteria. After incubation with antibiotics, cells were washed and lysed as detailed. Cell lysis mixture was serially diluted and plated on blood agar plates.

### Visualization of infected oral keratinocytes

Normal oral keratinocytes (NOK) cells were seeded and grown overnight on glass coverslips. Prior to infection, *P. gingivalis* strains (W83, V3128, and V3128C) were labeled with 2 µM BCECF-AM (ThermoFisher) for 30 minutes in PBS and washed. After labelling, NOK cells were infected at an MOI of 100:1 for 1 hour in an anaerobic chamber at 37°C. After infection, cells were washed 3 times with PBS and fixed using 4% paraformaldehyde. Actin was stained using Alexafluor 568 phalloidin (ThermoFisher). Coverslips were mounted using prolong anti-fade gold with DAPI (ThermoFisher). Slides were imaged on a Keyence BZ-X810 fluorescent microscope.

### Flow Cytometry

*P. gingivalis* was labeled with BCECF-AM and used to infect NOKs as detailed above. After washing, NOKs were trypsinized and suspended in flow cytometry buffer (PBS + 2% FBS). Cells were immediately run on a BD FACSymphony A1 flow cytometer, and the levels of BCECF-AM labeled bacteria associated with NOKs was assessed via FITC channel. Before sample injection, 0.5% trypan blue was added to quinch any extra-cellular fluorescence.

To measure *P. gingivalis* extra-cellular polysaccharide levels – bacteria grown on fresh blood plates were suspended in HBSS (Hanks balanced salts solution) and the optical density was adjusted to 0.5 OD_600_. To 1mL resuspended cells, 100 µg/mL of Concavalin A-FITC conjugated, 100 µg/mL wheat germ agglutinin-FITC conjugated, and 1 µM of Syto17 nucleic acid dye was added and incubated at room temperature for 1 hour under anaerobic conditions. After incubation, cells were washed 3 times with HBSS and run on a BD FACSymphony A1 Flow Cytometer. To distinguish bacteria from and cell debris, bacteria were gated for Syto17 (DNA binding dye) and the levels of extra-cellular polysaccharide capsule were assessed via FITC levels. A sample gating strategy is demonstrated in Fig. S7.

### Structure illumination microscopy

High precision glass cover slips were coated with poly-D-lysine (PDK) overnight and washed. From fresh plates, *P. gingivalis* strains were suspended in PBS to an OD of 1.0 and labeled with Hoechst dye (10 µg/mL) for 45 min at room temperature under anaerobic conditions. Labeled cells were washed with PBS and allowed to adhere to PDK coated coverslips for 30 min at room temperature. Excess liquid and unadhered cells were aspirated and cover slips were fixed using 4% paraformaldehyde for 15 minutes. Fixed cover slips were washed with PBS and incubated with 100 µg/mL Concavalin A-FITC conjugate (ThermoFisher) and 100 µg/mL wheat germ agglutinin-FITC conjugate (ThermoFisher) for 1 hour at room temperature. After incubation, cover slips were washed 3 times with PBS and mounted onto slides using Prolong Glass Antifade mountant. The next day sides were imaged on a Zeiss Elyra 7 structure illumination microscope.

### Transmission electron microscopy

*P. gingivalis* grown to stationary phase were centrifuged and washed in PBS. Cells were fixed at room temperature in 0.1 mol/L cacodylic acid buffer (pH 7) containing 3.6% (vol/vol) glutaraldehyde. Following fixation, cells were washed before secondary staining using 2% osmium tetroxide. Grids were visualized on a Zeiss EVO scanning electron microscope at the VCU microscopy core.

### Iron content assay

The levels of total iron in were measured using ferrozine via the Iron assay kit (Sigma). *P. gingivalis* strains were grown overnight in BHI broth. Overnight cultures were adjusted to an OD_600_ of 1.5. A 1.5 ml aliquot of culture was taken and washed twice using chelex treated HBS (25mM Hepes, 150 mM NaCl, pH 7.4). Cells were lysed in HBS + 0.5% SDS and incubated at 70°C for 10 minutes to fully denature proteins. After incubation, genomic DNA was sheared using a bath sonicator. Iron concentrations were then determined using the iron assay kit as per manufacturer instructions. The levels of iron were then normalized to the total protein content of each sample as measured via BCA assay.

### Bacterial adherence to hydrocarbons (BATH) assay

Overnight cultures grown in BHI were spun down, washed, and resuspended in PUM buffer (100 mM K_2_HPO_4_, 50mM KH_2_PO_4_, 30mM Urea, 1mM MgSO_4_) at an OD_600_ of 0.4. To 1.2 mL aliquots of cells, 0, 75, or 150 µL of N-hexadecane (Sigma) was added to the cells and vortexed vigorously for 2 minutes. After vortexing, mixtures were allowed to settle for 10 minutes and the OD_600_ of cells remaining in the aqueous phase was measured.

### Metal content assay

The cellular content of metals in *P. gingivalis* strains was assessed using inductively coupled plasma mass spectrometry (ICP-MS) analysis. W83 and V3128 bacterial cells from blood and BHI agar plates were suspended in 4 mls of BHI to obtain an OD_660_ of 1 - 1.4. Cells were pelleted by centrifugation at 7500 rpm for 10 minutes, washed twice with 4 mls of Chelex Buffer (0.05 M HEPES, 0.05 M NaCl), and suspended in 3 mls Chelex Buffer with 8 M Urea. The mixture was incubated for 1 hour at room temperature and bacterial cells were lysed using the sonication. Following centrifugation at 14,800 rpm for 30 minutes the supernatant was collected and used for ICP-MS analysis to determine metal content.

### Proteomics

#### Whole-cell proteomics

##### i) Sample preparation

Bacterial cells were suspended in a lysis buffer composed of 8 M urea/50 mM tetraethyl ammonium bicarbonate. Released proteins were digested with enzymes used: LysC (LysC to protein ratio of 1:100) and trypsin (trypsin to protein ratio of 1:100). The samples were then labeled with 10-plex TMT. Finally, samples were fractionated (12 fractions) using the basic pH RPLC method.

##### ii) Mass spectrometry analysis

Samples were analyzed on the Orbitrap Fusion Lumos (Thermo Scientific) with an on-line liquid chromatography: easy nLC 1200 (Thermo) with a 2 h running time per fraction. MS1 resolution was 120,000 and MS2 resolution was 30,000. HCD fragmentation method was used and collision energy for MS2 was 32.

##### iii) Database Search

The Proteome Discoverer 2.1 software was used as our database search tool. The *Porphyromonas gingivalis* (strain W83) UniProt (released on Apr 2017) with 2 missed cleavages allowed. Cleavage enzyme: trypsin, minimum AA length: 6, minimum peptides for protein: 1, fixed modification: Carbamidomethyl on cysteine residue and TMT on Lys and peptide N-terminal, dynamic modifications: Oxidation on M, and acetyl on N-terminus. 1% false discovery rate was applied for both peptide and protein levels

##### iv) Statistical analysis

p-value: calculated by Student’s t-test. q-value: calculated by significance analysis of microarrays (SAM) and permutation based false discovery rate (FDR). s0 value for SAM: 0.1 Reference for SAM ^14 15^. q-value is a kind of a modified p value but more stringent counting in multiple hypotheses. For q-value calculation, both p value and fold change were considered. Usually, proteins with q-value lower than 0.05 are considered to be statistically significant.

### Bioinformatics analysis

Sequence alignments of TonB related proteins were accomplished using ClustalW. Phyloanalysis and tree creation of TonB proteins from different species was created using MEGA12^16^. The genome region comparison using different bacteria was created using BioCyc^17 18^. Finally, an atomic model of the PG0785 TonB was generated using the AlphaFold3^19 20^.

## Results

### Bioinformatics analysis of PG0785

The *P. gingivalis* W83 genome encodes a putative TonB with ID PG0785 (new locus tag is PG_RS03445), a 693 bp gene coding for a 230 AA protein. Analysis of the PG0785 amino acid sequence revealed the highest similarity to TonB present on genomes of bacteria belonging to the Cytophaga-Flavobacter-Bacteroides (CFB) phylum (Fig. S1B). We performed a phylogenetic analysis of TonB members of the bacteroidales phylum with other well characterized TonB proteins found in *Escherichia, Helicobacter,* and *Vibrio.* These results reveal that *P. gingivalis* TonB clusters with *B. fragilis* TonB3, *B. thetaiotaomicron* TonB 4, and a TonB found in *T. forsythia* (Fig. 1A). Interestingly, these specific TonB proteins found in *B. fragilis* and *B. thetaiotaomicron* have been demonstrated to specialize in starch and polysaccharide utilization, interplaying with the starch utilization complexes (SusD/SusC) to transport carbohydrates across the membrane ^22 21 23^. A BLASTP comparison of PG0785-encoded protein revealed similarity to the C-terminal domain region of the *B. fragilis* 638R TonB3^21^ (Fig. S1A). *B. fragilis* TonB and the putative *P.gingivalis* TonB share 45% identity and 63% similarity (Fig. S1A). Furthermore, there was a high sequence identity observed in members of the *Porphyromonas* genus *(P. gulae* -93 % identity*, P. loveana* - 87.8% identity), but this dropped significantly for other members of the CFB bacteria indicating that the PG0785 TonB protein possesses features unique to the genus *Porphyromonas* (Fig. S1B). Very high conservation in the gene content was observed when comparing the TonB loci in *P. gingivalis* W83 and in the *B. fragilis* 638R indicating that the entire locus composed of a TonB and the accessory proteins ExbBD is conserved (Fig. 1B) ^21^. Furthermore, similar locus is also present in *Bacteroides thetaiotaomicron* KPPR-3 NZ and *Tannerella forsythia* 920AC (Fig. 1B). While the *B. fragilis* TonB3 has been characterized, the *B. thetaiotaomicron* and the *T. forsythia* loci are yet to be examined. The locus is composed of seven genes with lower similarity for the TonB protein (> 40%) but higher for the ExbB and ExbD proteins (>50% identity). Of note is the finding that there are 2 ExbD proteins, while one of the ExbD (ExbD2) is from TolR family, and the other (ExbD) is from the TonB lineage.

**Figure 1.**
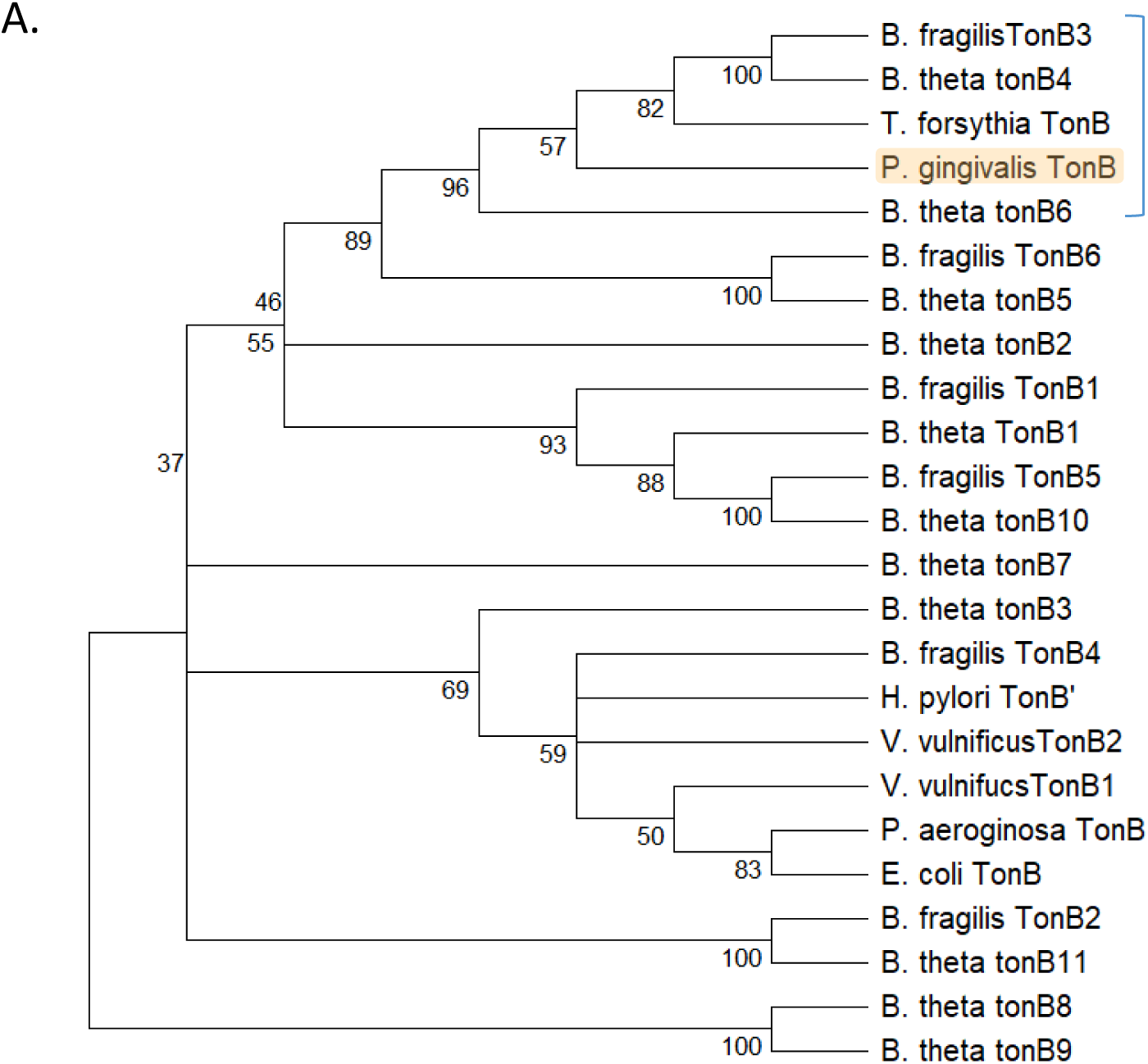
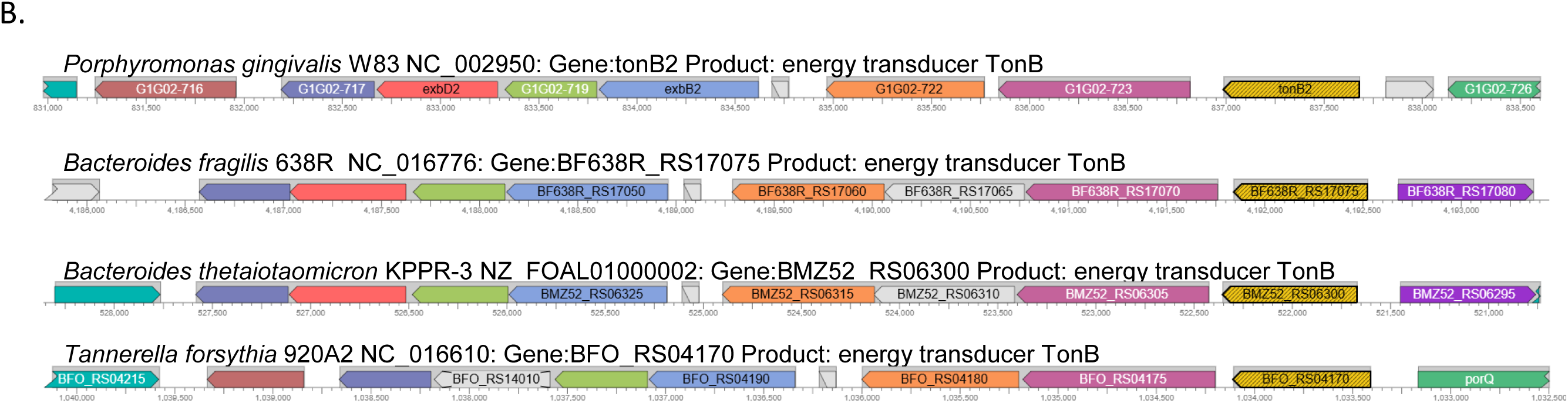
Comparison of PG0785 with proteins present in Bacteroidota. A. Phylo- analysis of related TonB proteins inferred using the Neighbor-Joining method. The optimal tree with the sum of branch length = 15.484 is shown. The percentage of replicate trees in which the associated taxa clustered together in the bootstrap test (500 replicates) are shown below the branches. The *P. gingivalis* TonB encoded by PG0785 is shown as shaded in orange. B. The genome region comparison using different bacteria was created using BioCyc. PG0785 in *P. gingivalis* W83 is shown as yellow arrow (tonB2), the corresponding TonB3 from *B. fragilis* 638R (BF_RS17075) and TonBs from *B. thetaiotaomicron* KPPR-3 NZ (BMZ52_RS06300) and *T. forsythia* 920A2 NC (BFO_RS04170) are also shown as yellow arrows.

Despite differences in the full length amino acid sequences, the C-terminal domain of *P. gingivalis* TonB reveals moderate sequence similarity to TonBs with solved structures: *Helicobacter pylori* ^24^ (6sly.1.A) (O25899, 41.2% identity)*, E. coli*, and *Pseudomonas aeruginosa* (Q05613.1, 33.8% identity) (Fig. 2A) ^25 26^. An AlphaFold model has been generated for the full length of the *P. gingivalis* TonB composed of the transmembrane domain (TM domain), proline-rich low complexity region (LCCR domain), and C-terminal domain (CTD domain) (Fig. 2B-C). There was a high conservation of amino acids that are important in the function of TonB proteins. The YP motif, the most conserved feature in TonB proteins, is present in PG0785 TonB at residues 14-15 (Fig. 2A, residues 23-24 in the alignment). The tyrosine residues have been demonstrated to interact directly with TBDT in *E. coli* and *P. aeruginosa,* and mutation of either Y or P leads to loss of functionality of the TonB^27^. Also, other conserved residues: A, QG, V and DG were identified in the *P. gingivalis* TonB (Fig. 2A). The last beta-strand of the C-terminal domain is implicated in directly binding to the Ton Box in TBDT (beta-strand 3), an interaction that is key to supplying the proton motive force allowing for transport of molecules. Notably, there is a high degree of conservation in the final beta-strand of *P. gingivalis* with *B. fragilis* TonB3 and *B. thetaiotaomicron* TonB4 that is different from more distantly related TonB orthologues in *E. coli* and *P. aeruginosa.* This conservation implies a possible compatibility between *Bacteroides* and *Porphyromonas* TonB-dependent transporter orthologues.

**Figure 2.**
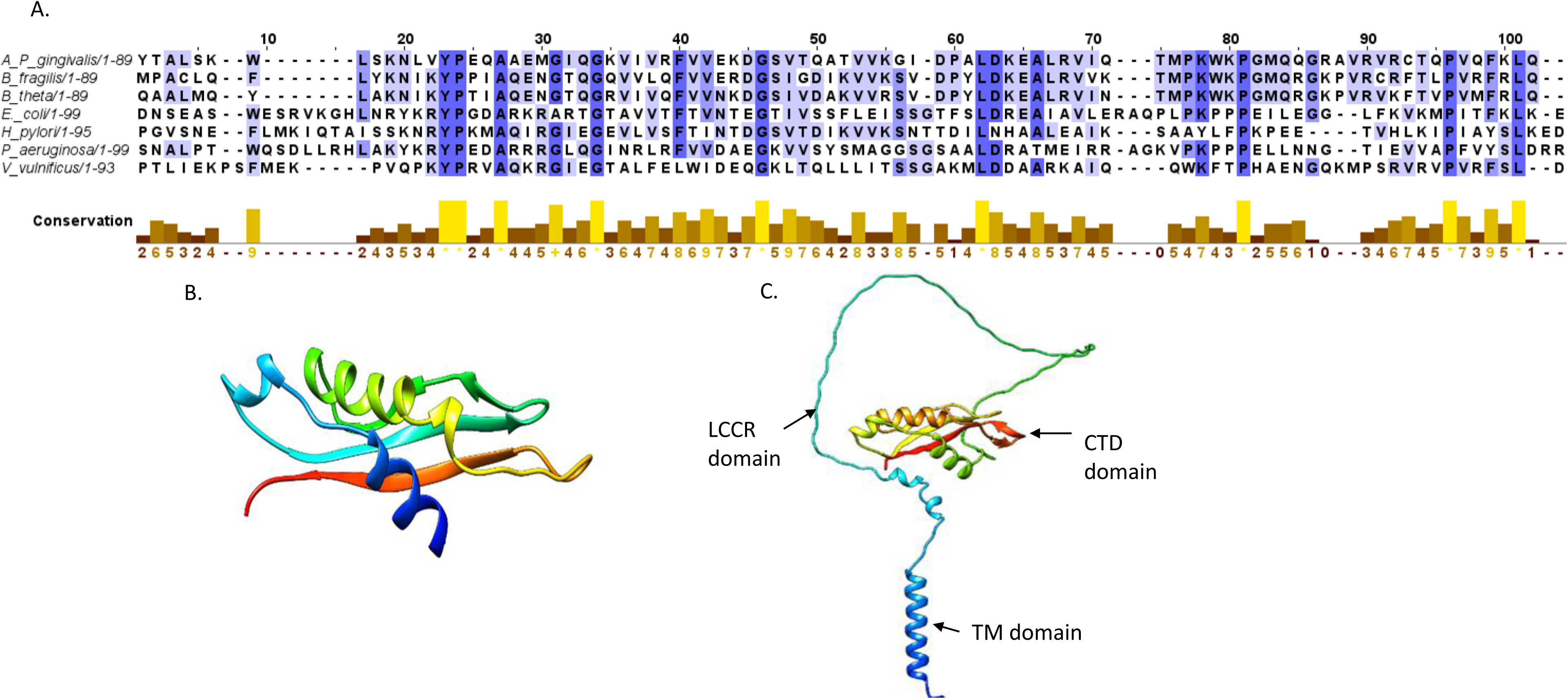
Structural characteristics of the *P. gingivalis* W83 TonB encoded by PG0785. A. Alignment of *P. gingivalis* TonB (PG0785) CTD to proteins with known structures present in SwissProt. Residue conservation for each position is scaled from 0 (no conservation) to 10 (represented as a *, complete conservation) and shown below alignments. B. Model of a template alignment of an NMR solution structure of *Helicobacter pylori* TonB – CTD domain with a predicted structure of the PG0785 protein. 30.59% identity was detected between the proteins. C. An AlphaFold generated model of the PG0785 protein. The three regions of the protein are shown: transmembrane domain (TM domain), proline-rich low complexity region (LCCR domain), and C-terminal domain (CTD domain).

### TonB is required for pigment accumulation

In order to be able to determine the role of TonB in *P. gingivalis* we generated an insertion mutant in the PG0785 gene (*tonB*). An *ermF-ermAM* cassette^11^ was inserted at the *Nru*I site of *tonB* (located at the 82bp of the gene) thus generating mutant designated V3128. The mutation was complemented by providing an intact copy of the *tonB* on the pG108 vector and the strain was designated V3128 complemented (V3128C). While the V3128 grew well on blood agar plates it did not form the black pigmentation as observed in the parental W83 strains (Fig. 3A). Complementation via placement of the PG0785 gene on a plasmid restored black pigmentation. The TonB-deficient mutant (V3128) grew on blood and BHI agar plates but appeared compromised in broth culture. There was no significant difference in growth of the initial broth inoculum (P1), however, subsequent passages in broth media saw a significant drop in growth (P2 and P3) (Fig. S2). This growth deficiency was restored in the V3128C strain, which could be indefinitely passaged in broth media, similar to the wild type.

**Figure 3.**
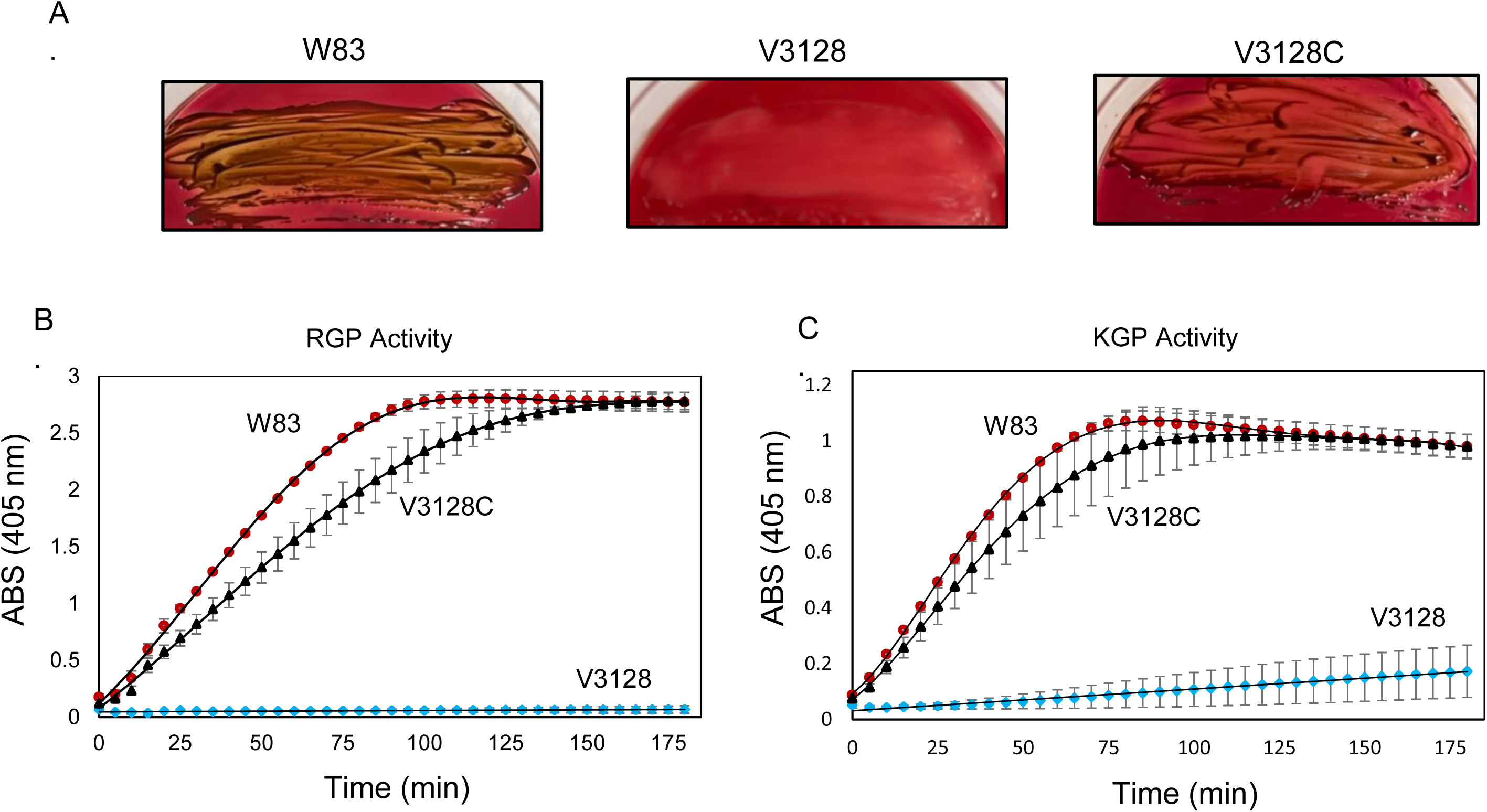
*Porphyromonas gingivalis* TonB is required for pigmentation on blood agar plates and gingipain activity. A. Wild Type W83, V3128 *tonB* mutant, and V3128 TonB-complemented *P. gingivalis* were grown on TSA 5% sheep blood agar plates for approximately 7 days and imaged. B.-C. Protease assay of bacteria grown in BHI broth overnight. The whole cell and supernatant protease activity of Arg-gingipain and Lys-gingipain was chromogenically measured over 180 minutes using 1mM BApNA (RGP) or 0.5mM TGPLpNA (KGP). The x-axis represents the time in minutes of the measurement’s recordings. The y-axis represents the activity of proteases as assessed by measuring absorbance at 405 nm.

### Loss of TonB reduces gingipain activity

Gingipain activity was demonstrated to play a role in pigment accumulation in *P. gingivalis*^3 7 28^. The enzyme activity rates for both Arg and Lys gingipain activities were determined for the wild type (W83), V3128, and V3128C strains using BApNA (1mM) and TGLpNA (0.5mM). Both, whole cell and culture supernatant (secreted) protease activities were measured. The whole cell Arg gingipain activity for WT was 13.5 µmol/min (Fig. 3B). The V3128 strain had no Arg gingipain activity. Complementation restored the activity to 10.1 µmol/min (Fig. 3B). These results were also mirrored in the whole cell Lys gingipain activity: WT activity was 7.2 µmol/min; the V3128 strain had no Lys gingipain activity, and the V3128 complement one had 6.2 µmol/min (Fig. 3C).

Similar results were seen with the secreted protease activity. The WT secreted gingipain activity was measured at 10.1 µmol/min (RGP) and 4.5 (KGP); There was no secreted gingipain activity in the V3128 TonB deficient strain (Fig. S3A-B). Complementation restored gingipain activity to 4.7 µmol/min (RGP) and 3.1 µmol/min (KGP) (Fig. S3A-B).

### Significant alterations in *P. gingivalis* proteome are observed in the absence of TonB

SDS-PAGE analysis showed drastic changes in protein profile resulting from the TonB deficiency (Supplemental Fig. 5). To gain an insight into the protein profile changes of the bacterial cells we performed whole cell proteomics on the WT and V3128 TonB-deficient strains. We examined proteome profiles from bacterial cells grown on both blood agar and BHI agar plates. Three biological replicates were prepared for samples grown on blood agar plates and two biological replicates were examined for bacteria grown on BHI agar plates. In total, 1223 *P. gingivalis* proteins were identified (Supplemental Table 1). Comparison of *P. gingivalis* strains V3128 to W83 protein abundance of bacterial cells derived from blood agar plates showed that 301 proteins were significantly regulated (q <0.01) (Fig. 4A, Supplemental Table 2). The top regulated proteins are shown in Tables 1 and 2. Upregulated were proteins PG1127 (AsnC, transcriptional regulator), PG1345 (glycosyl transferase), PG0627 (RNA-binding protein), PG1810 (2-oxoglutarate oxidoreductase), and several uncharacterized proteins. Also, upregulated were 50S and 30S ribosomal proteins. Among the downregulated ones were: PG0785 (TonB, thus verifying our mutagenesis data), PG0456 (PHP N-terminal domain protein), CydA (cytochrome d ubiquinol oxidase), PG1503 – 05 locus coding for LytB-related protein, NAD dependent protein, and radical SAM domain protein, PG0717 (putative lipoprotein), PG0174 (pyridine nucleotide – disulphide oxidoreductase), PG0437 (polysaccharide export protein), and PG0538 (outer membrane efflux protein). Several uncharacterized proteins were also detected. The PG0437 is the first gene of a locus also coding for a capsule synthesis-associated tyrosine kinase, Ptk1^29^. The LytB locus (PG1502-5) is involved in the methyl erythritol phosphate (MEP) pathway that is essential for bacterial viability ^30 31^. LytB, the last enzyme in that pathway, contains an iron-sulfur cluster [4Fe-4S] and is required for the last step of MEP catalysis^30 31^. Finally, the upregulated locus PG0538-541 encodes an efflux system composed of a TolC -like protein, periplasmic adaptor protein and AcrBDF-like protein (Supplemental Fig.6). The system, named XebCBA, was shown to pump out multiple substances such as rifampin, puromycin and ethidium bromide^32^.

**Figure 4.**
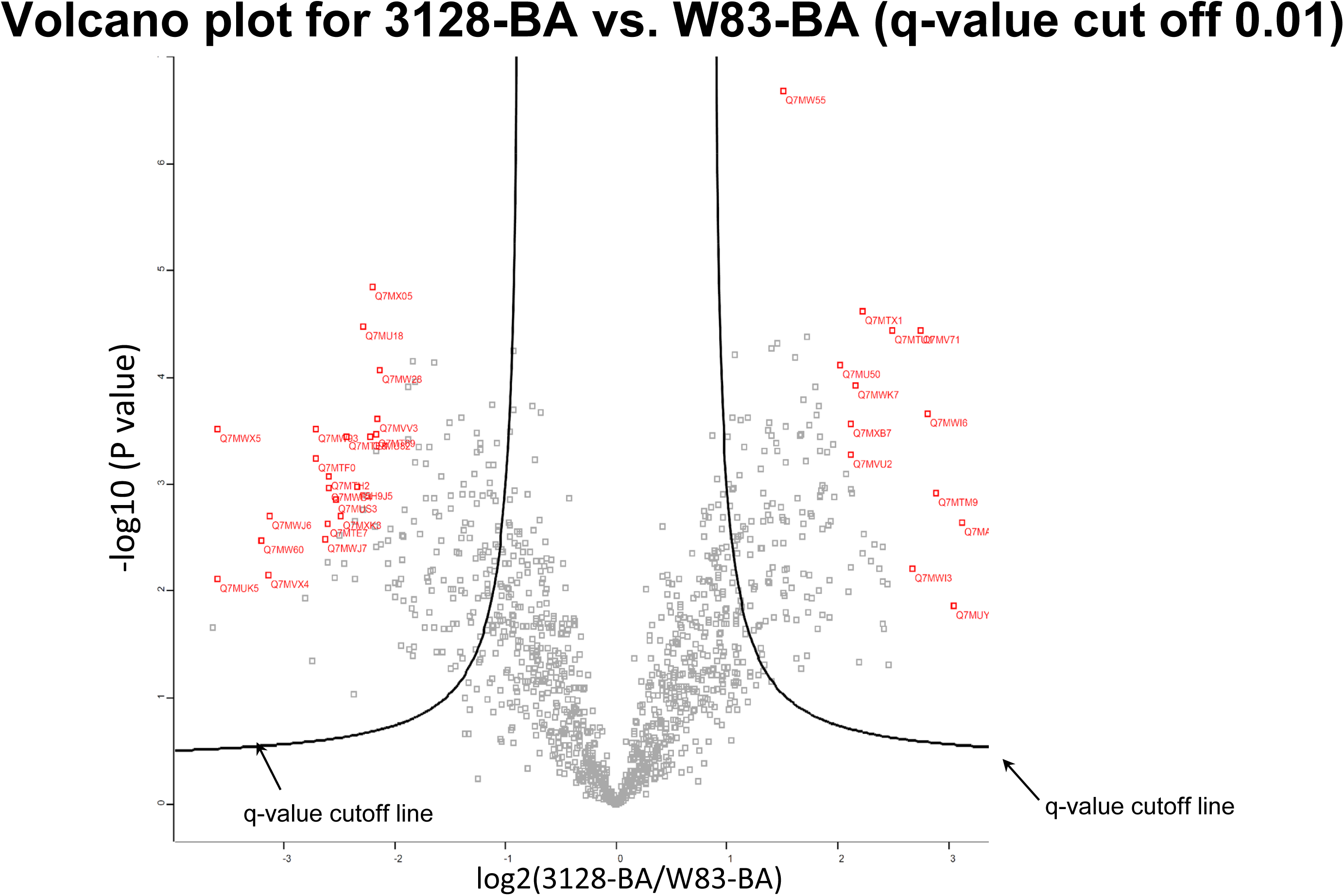
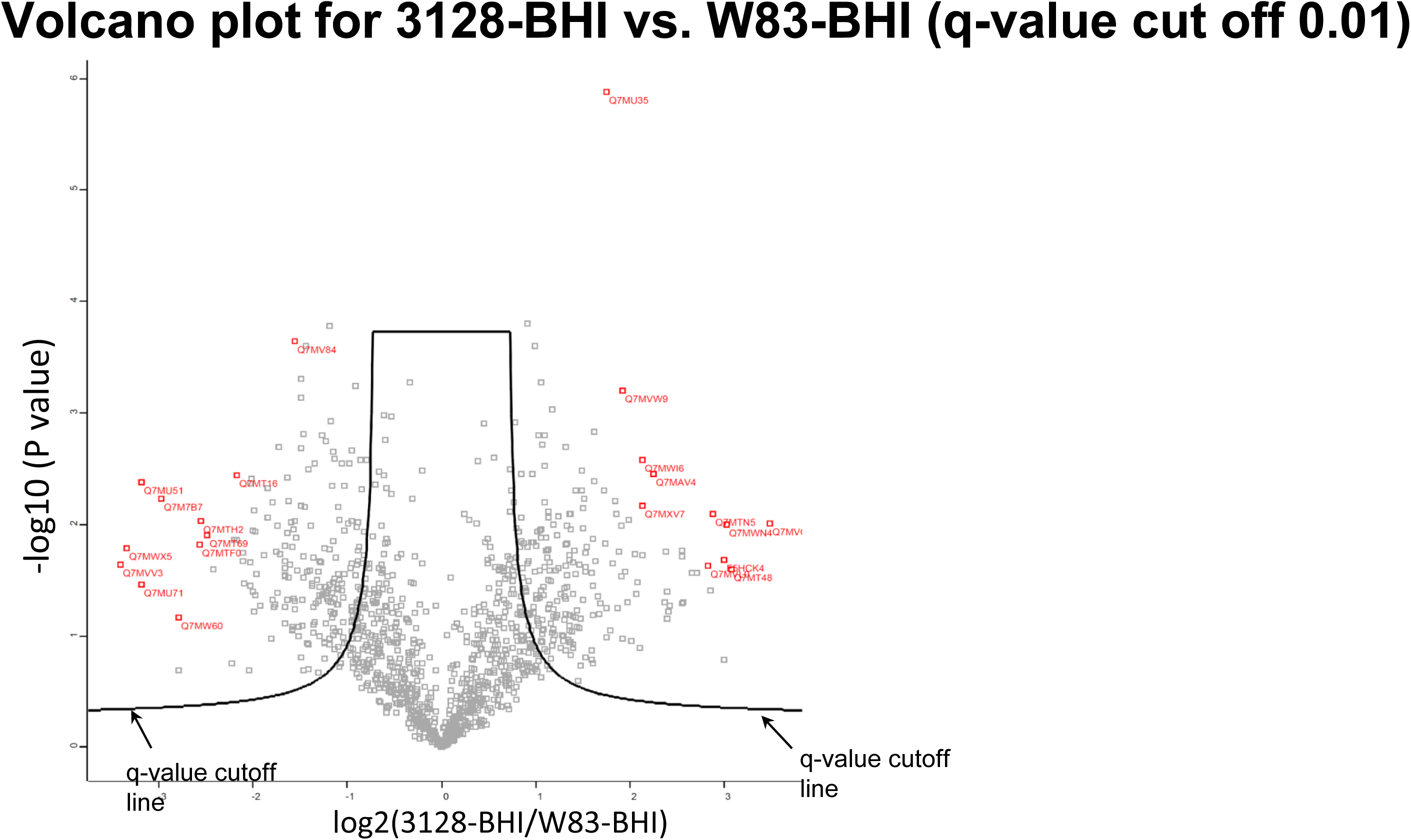
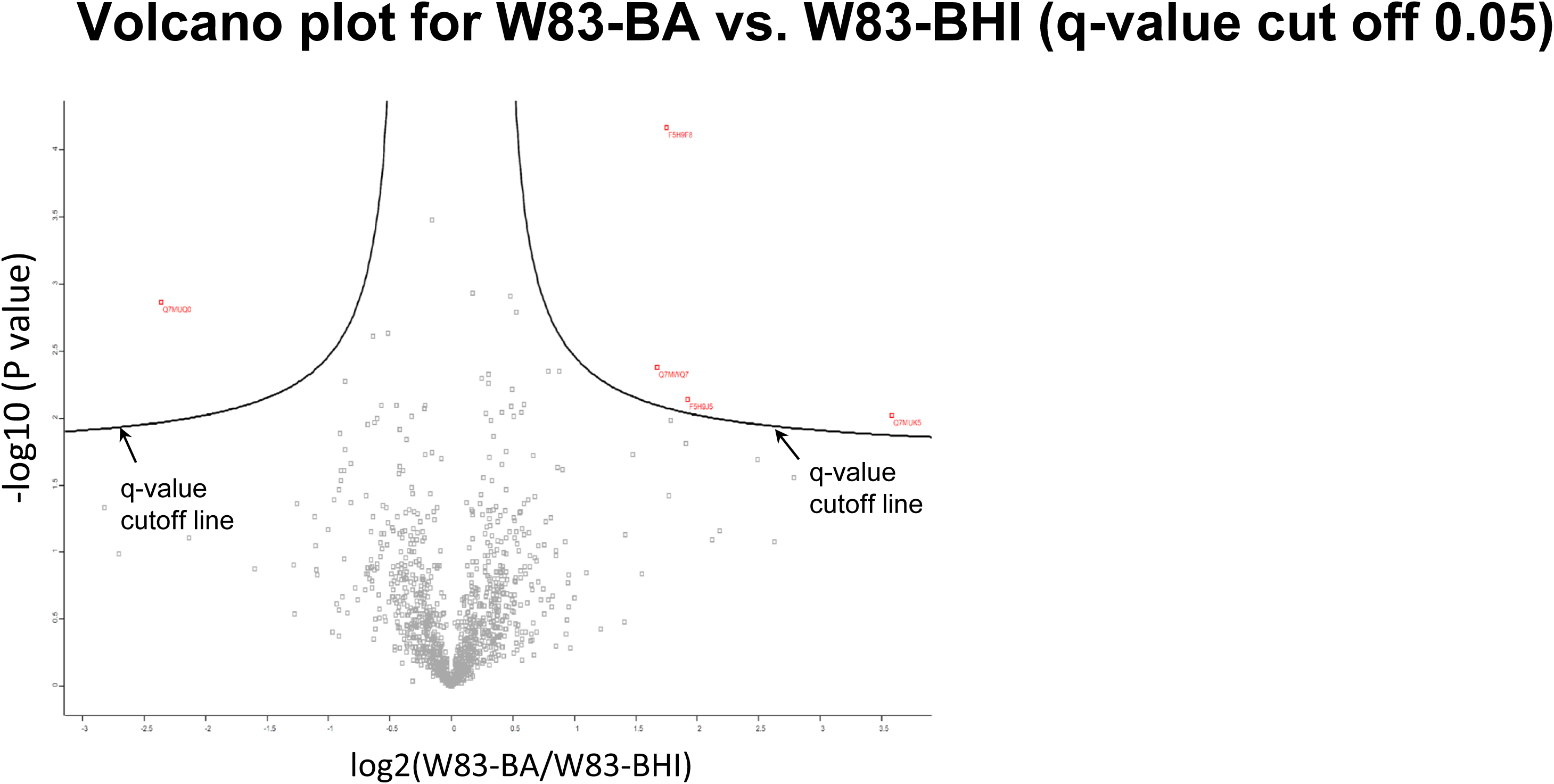
TonB deficiency alters the protein profile of *P. gingivalis* cells. A. Comparison of a protein profile of V3128 and W83 cell lysate proteins derived from bacteria grown on blood agar plates. The proteins outside the q-value cutoff line can be considered to be significant with 1% of possibility being false positive. Highlighted in red font are the accession numbers of the proteins with most high confidence. B. Comparison of a protein profile of V3128 and W83 cell lysate proteins derived from bacteria grown on BHI agar plates. The proteins outside the q-value cutoff line can be considered to be significant with 1% of possibility being false positive. Highlighted in red font are the accession numbers of the proteins with most high confidence. C. Effect of growth media on protein expression in *P. gingivalis* W83. Comparison of a protein profile of W83 cell lysate proteins derived from bacteria grown on blood agar plates to W83 cell lysate proteins from cells grown on BHI agar plates. The proteins outside the q-value cutoff line can be considered to be significant with 5% of possibility being false positive. Highlighted in red font are the accession numbers of the proteins with most high confidence.

**Table 1.**
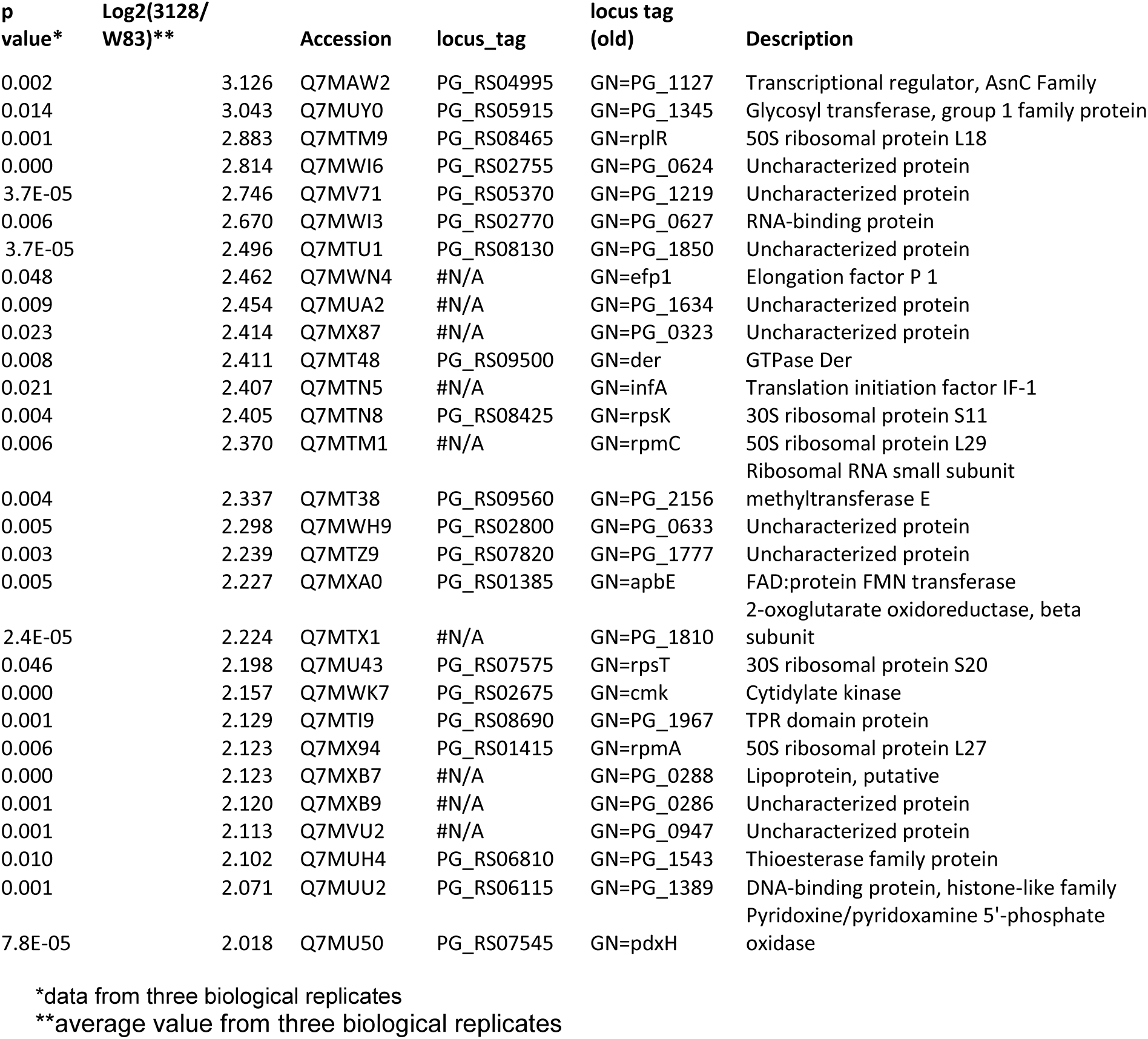
List of the most differentially upregulated proteins in TonB-deficient mutant V3128 when compared to parental W83 strain. Cell lysates from bacteria grown on blood agar plates were used for analysis.

**Table 2.**
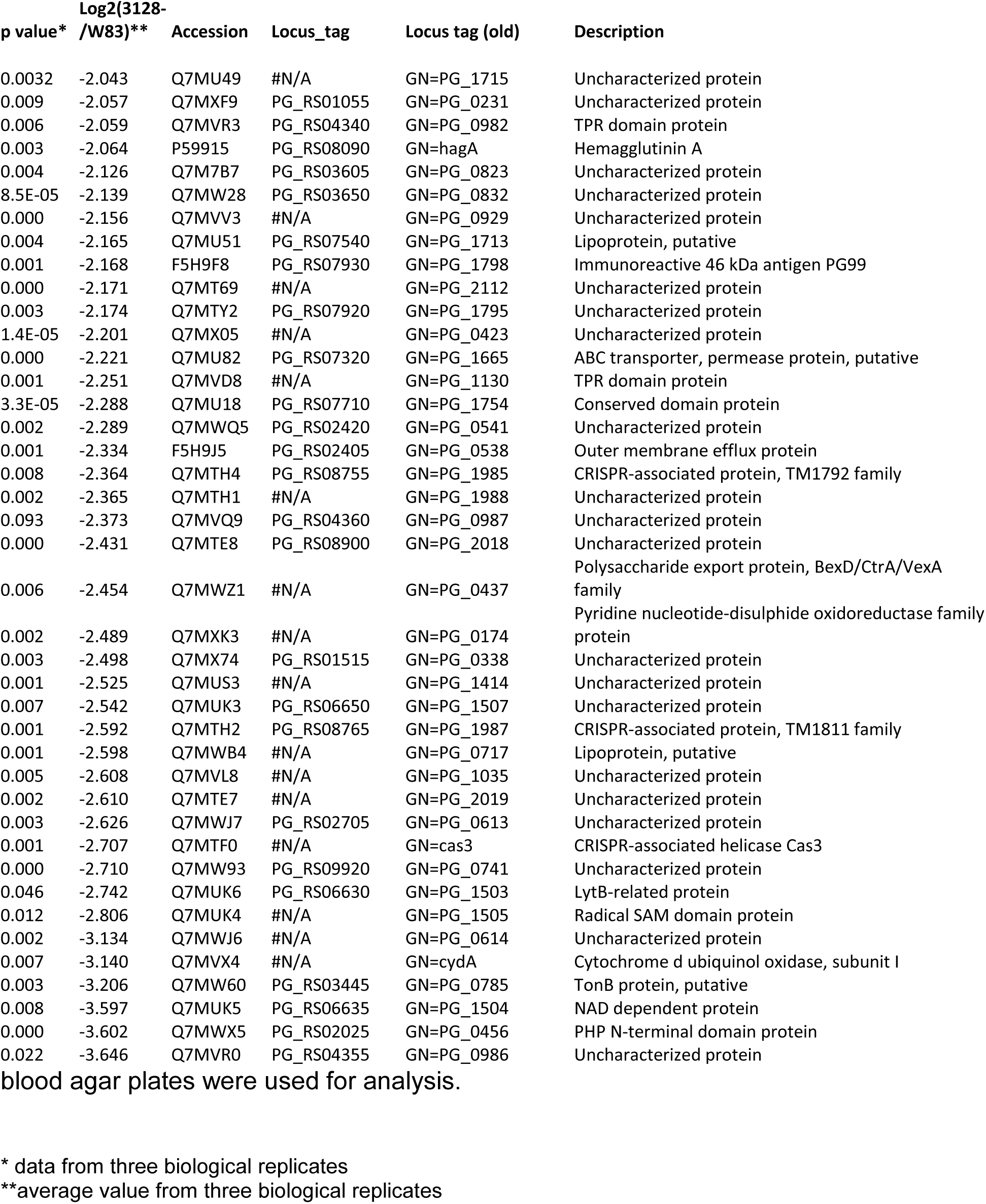
List of the most differentially downregulated proteins in TonB-deficient mutant V3128 when compared to parental W83 strain. Cell lysates from bacteria grown on blood agar plates were used for analysis.

**Table 3.**
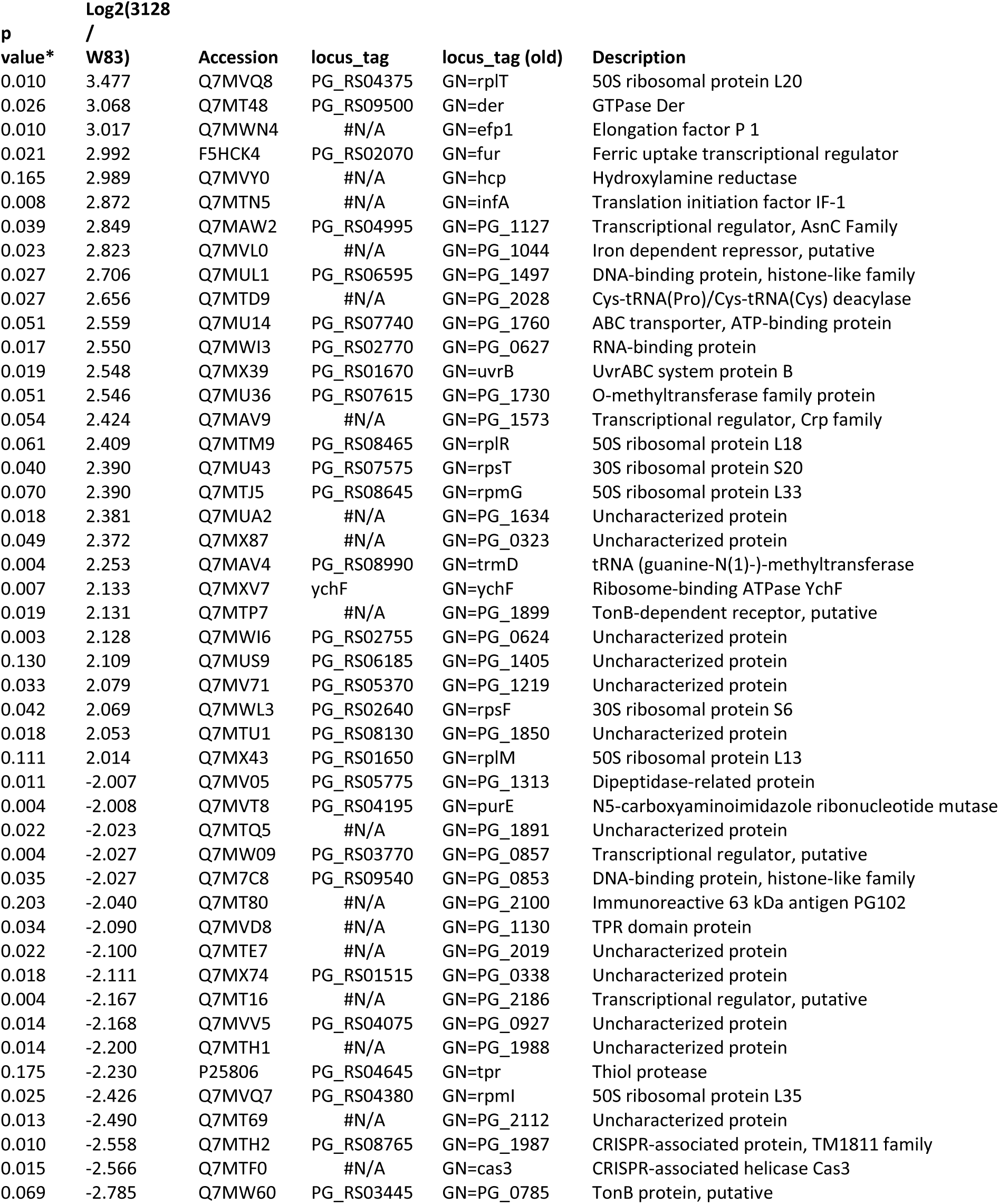
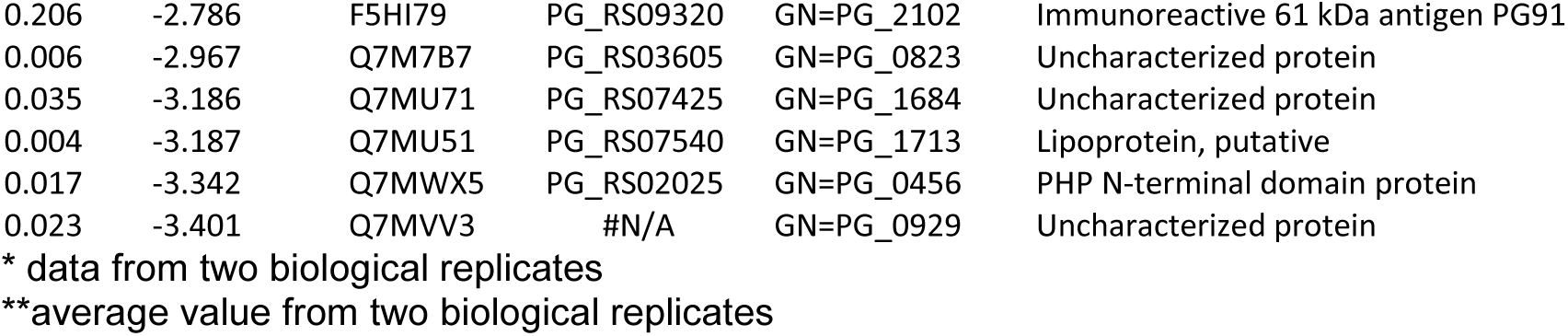
List of the most differentially regulated proteins in TonB-deficient mutant V3128 when compared to parental W83 strain. Cell lysates from bacteria grown on BHI agar plates were used for analysis.

**Table 4.**
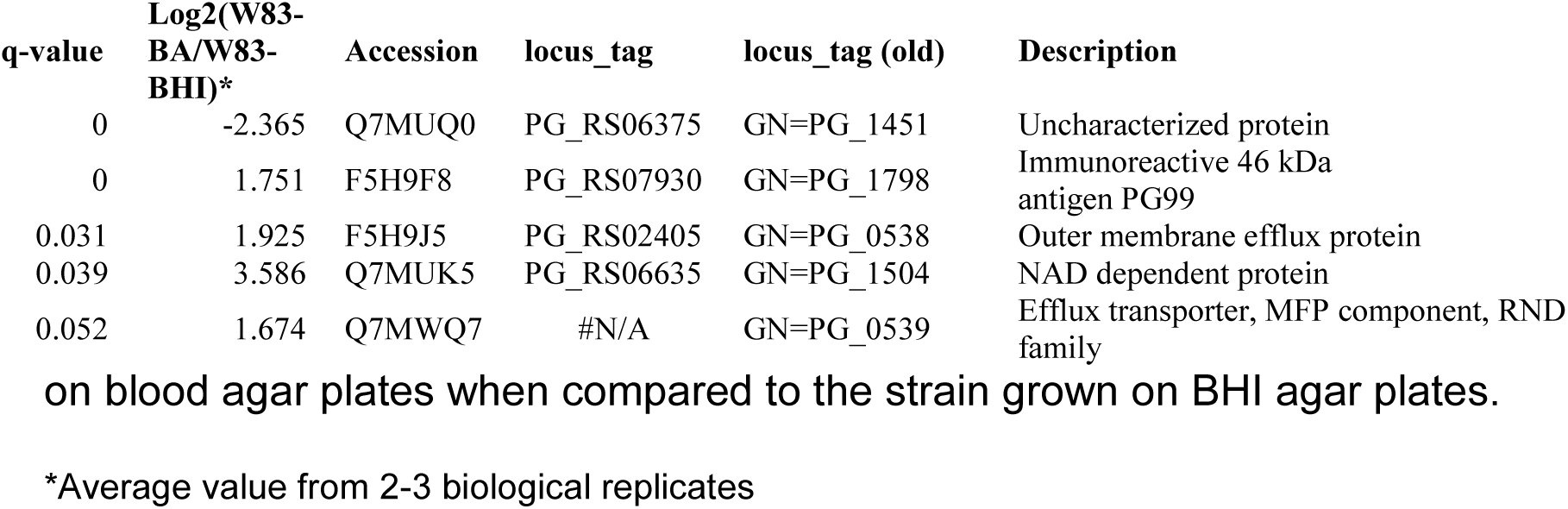
List of most differentially regulated proteins in the parental W83 strain grown on blood agar plates when compared to the strain grown on BHI agar plates.

| q-value | Log2(W83-BA/W83-BHI)* | Accession | locus_tag | locus_tag (old) | Description |
| --- | --- | --- | --- | --- | --- |
| 0 | -2.365 | Q7MUQ0 | PG_RS06375 | GN=PG_1451 | Uncharacterized protein |
| 0 | 1.751 | F5H9F8 | PG_RS07930 | GN=PG_1798 | Immunoreactive 46 kDa<br>antigen PG99 |
| 0.031 | 1.925 | F5H9J5 | PG_RS02405 | GN=PG_0538 | Outer membrane efflux protein |
| 0.039 | 3.586 | Q7MUK5 | PG_RS06635 | GN=PG_1504 | NAD dependent protein |
| 0.052 | 1.674 | Q7MWQ7 | #N/A | GN=PG_0539 | Efflux transporter, MFP component, RND<br>family |

\*Average value from 2-3 biological replicates

Comparison of protein profile of bacterial cells from the parental strain *P. gingivalis* W83 to the mutant V3128 grown on BHI plates has shown differential regulation of 423 proteins (q < 0.01) (Fig. 4B, Supplemental Table 3). The top most differentially regulated proteins are shown in Table 2. As with data derived from blood plates, upregulation of PG0627 (RNA binding protein) and PG1127 (AsnC transcriptional regulator) was observed, however, many other proteins were also regulated. Among those were: Hcp (putative hydroxylamine reductase/ nitric oxide reductase), PG1497 (DNA-binding protein, histone - like), PG1573 (Crp-like transcriptional regulator), GTPase der, and several uncharacterized proteins. Also, ribosomal proteins (30S and 50S) were regulated. The downregulated proteins in bacteria grown on BHI plates included PG0785 (TonB) and PG0456 (PHP N-terminal domain protein) that were also downregulated in bacteria grown on blood agar plates. However, several other proteins were also differentially downregulated: PG1713 (lipoprotein), Tpr (thiol protease), transcriptional regulators (PG2186, PG0857), PG0853 (histone-like DNA-binding protein), PG2100 (immunoreactive 63kDa antigen), PG1987-8 (CRISPR – system proteins). Finally, several uncharacterized proteins were downregulated.

While comparing the control samples, *P. gingivalis* parental W83 strain grown on blood agar plates to that grown on BHI agar plates has identified 5 differentially regulated proteins (q <0.05)(Fig. 4C), no significantly differentially regulated proteins were found for the *P. gingivalis* TonB-deficient mutant V3128.

The above data shows extensive proteome modification in the absence of TonB. Although the TonB protein has been shown to play a role in energizing transport of metals through the TonB-dependent receptors, our data shows that the TonB plays a wider role in *P. gingivalis*.

### Absence of TonB alters metal content profile in *P. gingivalis* cells

Classically, the role of TonB in iron metabolism has long been established, playing a key role providing the energy necessary to transport iron bound siderophores across the bacterial cell membrane, most effectively demonstrated in *E. coli* and other siderophilic bacteria such as *Vibrio* species. Thus, to gain insight into the role TonB may be playing in iron metabolism in *P. gingivalis* we measured the total iron levels in the bacteria using a ferrozine assay (Fig. 5A). Unexpectedly, when grown on BHI we found no difference in the levels of total iron in the WT, V3128, or V3128C.

**Figure 5.**
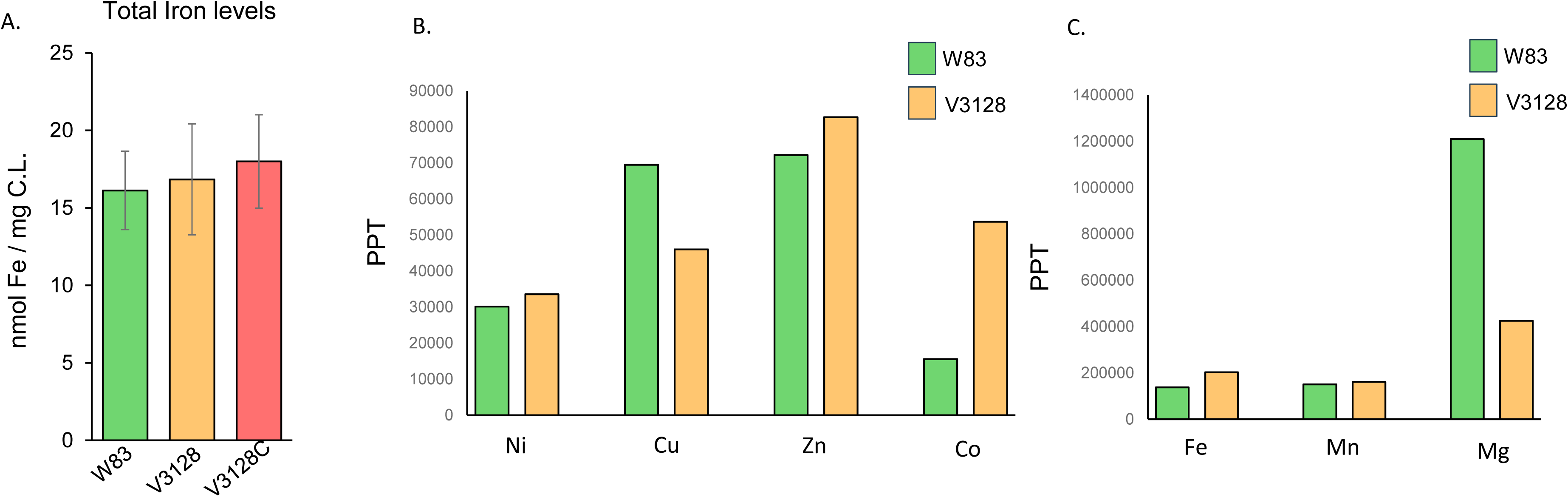
*P. gingivalis* TonB affects metal content of copper, cobalt, and magnesium but not iron. A. The levels of total iron in WT, V3128, and V3128C strains grown in BHI media were measured using a ferrozine based assay. The levels of iron were normalized to the total amount of cell lysate as measured by BCA assay. B-C. WT and V3128 TonB-deficient strain grown on BHI were analyzed using an ICP-MS to determine the amount of iron (Fe), magnesium (Mg), manganese (Mn), zinc (Zn), nickel (Ni), copper (Cu) and cobalt (Co) in each sample. Metal content results for Ni, Cu, Zn and Co are shown in Panel B and results for Fe, Mg and Mn are shown in Panel C.

To confirm these results and to comprehensively analyze the metallome, we performed ICP-MS on the WT and V3128 strains grown on BHI agar plates. Strains were analyzed for the levels of nickel, copper, zinc, cobalt, iron, manganese, and magnesium (Fig 5B-C). As with our ferrozine data, the levels of iron in both WT and V3128 were similar, revealing that Fe metabolism in the V3128 was not significantly altered. However, differences in other metals were noted. In the V3128 mutant strain, the levels of Zn and Cu were elevated and the levels of Co were decreased when compared to the WT. However, most drastic was changes in the Mg levels from 1.21 x 10^6^ PPT in the WT to 4.26 x 10^5^ PPT in V3128 strain.

In addition, the metallome of bacteria grown on TSA-blood agar plates were also analyzed (Fig. S4), and reveals similar trends in the Zn, Cu, Co, and Mg levels to those observed on BHI with the notable exception of Fe, which is orders of magnitude higher in the WT strain (at 3.4 x 10^6^ PPT) than in the V3128 TonB deficient strain (1.68 x 10^5^ PPT). When grown on blood agar plates, *P. gingivalis* coats its outer surface in heme giving it a distinct black color, a key phenotype that does not occur when grown on BHI agar plates. The V3128 does not turn black when grown on blood agar plates (Fig. 3A). Thus, the drastic differences in the iron levels derived from blood agar ICP-MS is a reflection of the loss of black pigmentation.

### TonB is required for survival of *P. gingivalis* with host cells

To determine if *P. gingivalis* TonB helps to mediate survival with host cells, we investigated the ability of the V3128 strain to survive with human immortalized keratinocyte cells as well as the HL60 cell line. As demonstrated in Fig. 6A, the wild type strain was recovered efficiently in both NOK and HL60 cells. There was significantly less V3128 recovered from NOK cell cultures and there was no *P. gingivalis* detected from HL60 cultures (Fig. 6B). Complementation restored the survival of the V3128C strain to wild type levels in NOK cell cultures (Fig. 6A).

**Figure 6.**
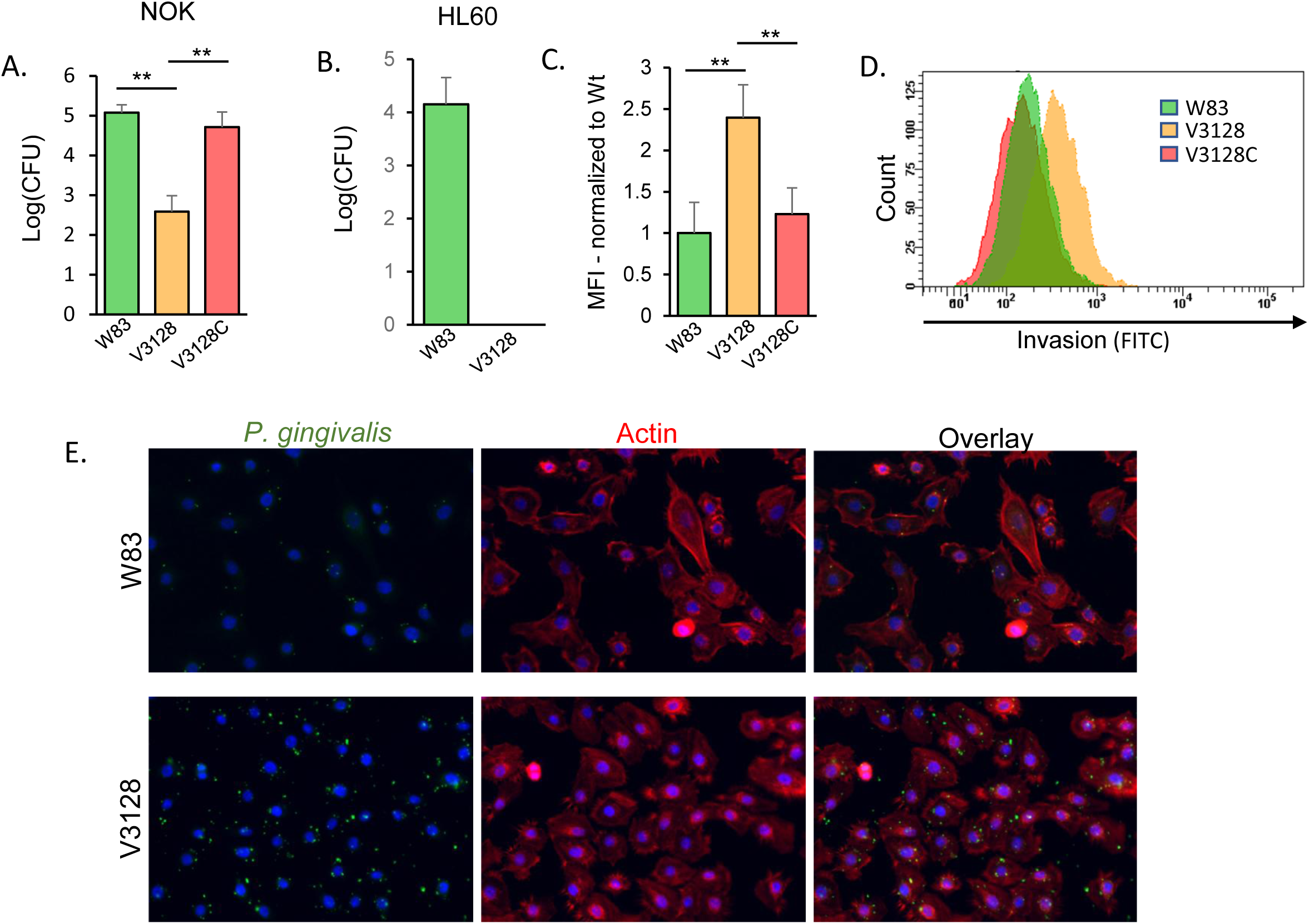
TonB is required for *P. gingivalis* survival following interaction with host cells but promotes microbial invasion. A.-B. Total interaction assay using normal oral keratinocytes (NOKs) (A) and HL60 (B) of WT W83, V3128 *tonB* mutant, and complemented strain (V3128C) infected at an MOI of 100:1. No viable bacteria were recovered for the V3128 strain with HL60s. C. Normal oral keratinocytes (NOKs) were infected with stained *P. gingivalis* strains at an MOI of 100:1 before flow cytometry. Data is represented as the median fluorescence intensity (MFI) of 10,000 cells normalized to the intensity of the WT W83 signal. Experiment was repeated in 3 biological replicates. D. Representative histogram of data expressed in panel C. E. Representative fluorescence microscopy images of normal oral keratinocytes infected with WT W83 and V3128 *tonB* mutant. *P. gingivalis* stained with BCECF-AM (green), actin was stained with phalliodin-AF568 (red), and nuclei are stained with DAPI (blue).

To determine if association or entry of the V3128 was compromised we performed confocal microscopy on NOK cells infected with the WT and V3128 strain. Unexpectedly, we observed a significant increase in the V3128 strain adhering to and invading cells when compared to the wild type (Fig. 6C). These results were quantified using flow cytometry to measure total invasion of host cells (Fig. 6C-D). There was approximately twice the amount of V3128 strain as wildtype strain in NOKs. Complementation restored the invasion/interaction efficiency back to wild type levels. Additionally, we observed no difference in the labeling efficiency of BCECF-AM between WT, V3128, and V3128C strains.

Overall, the results show that the absence of TonB severely impairs the ability of *P. gingivalis* to survive with host cells despite an observed increase in the invasion/interaction of the V3128 mutant strains when compared to the WT and complemented strains.

### Loss of TonB alters the surface of *P. gingivalis*

Proteomics experiments revealed that capsule associated with kinase Ptk1 was one of the most down-regulated proteins in the TonB knockout strain (Table 1). That kinase was shown to be indispensable for capsule production and in agreement with that we tested our mutant strain for the presence of the capsule ^33^. Using a combination of lectins that has previously been demonstrated to bind to the W83 capsule (wheat germ agglutin and concavalin-A), the capsule was imaged using structure illumination microscope (SIM). As can be seen, there was a near complete breakdown of the extra-cellular capsule formation in the absence of TonB when compared to the wild type (Fig. 7A). Capsule formation was restored when the TonB deficient strain was complemented (Fig. 7A). Using flow-cytometry, we quantified this change and saw there was significant decrease in the extra-cellular polysaccharides that make up the capsule in the V3128 TonB-deficient strain (Fig. 7B).

**Figure 7.**
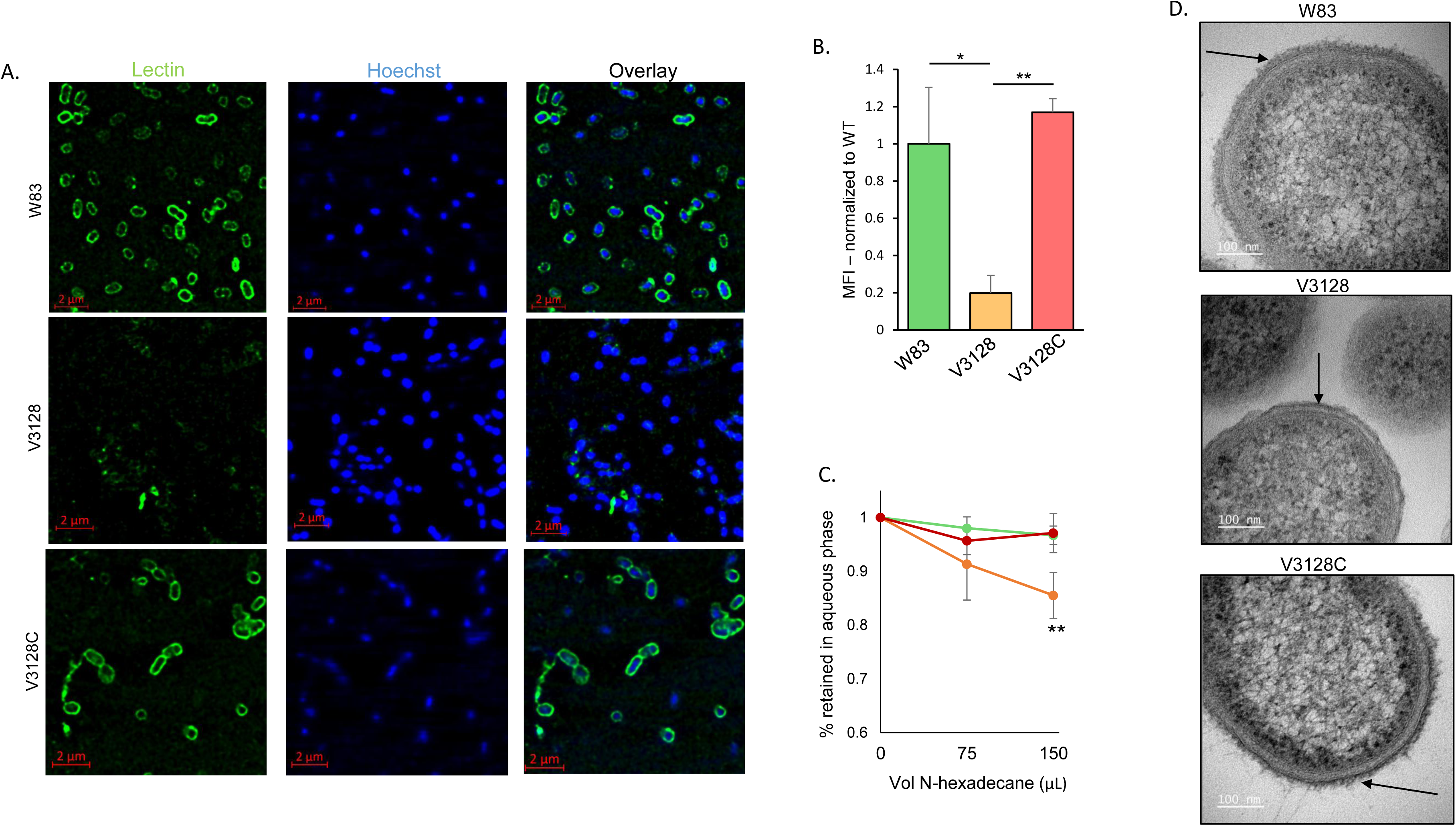
TonB is necessary for extra-cellular capsule polysaccharide formation. A. *P. gingivalis* strains were labeled with Hoechst dye then seeded and fixed onto poly-D-Lysine coated cover slips. A combination of FITC-conjugated Wheat Germ Agglutin (WGA) lectin and Concavalin-A (ConcA) lectin was used to stain the capsule. Images were taken on a Zeiss Elyra 7 structure illumination microscope (SIM). B. Transmission Electron Microscopy of *P. gingivalis* strains. Arrow indicate presence of surface capsule in the Wildtype and complement strains versus the reduced capsule in the V3128 *tonB* mutant strain. Panel C. *P. gingivalis* strains were stained with a mixture of FITC conjugated WGA and ConcA lectins. Bacteria were washed and the median fluorescence intensity was measured on a FACSymphony A1 flow cytometer. Data is representative of the mean fluorescence intensity of the 3 biological replicates normalized to the signal of WT W83. D. Images of *P. gingivalis* strains were taken use transmission electron microscopy (TEM). The presence of the electron dense region of WT and V3128C strains is highlighted by arrows.

Changes in extra-cellular polysaccharide can alter the hydrophilic or hydrophobic nature of the membrane. A loss in a (hydrophilic) capsule would possibly make the outer-membrane more hydrophobic in nature. We tested the ability of the V3128 to adhere / interact with hydrocarbons using N-hexadecane. The outer-membrane of *P. gingivalis* W83 is typically very hydrophilic due to the presence of a capsule and anionic-LPS in the cell membrane, thus, the WT and complement strains do not adhere or interact with hydrophobic N-hexadecane and fully remain in the hydrophilic phase (Fig. 7C). However, a significant portion of the TonB mutant strain did end up in the hydrophobic phase (approximately 17%), indicating a change in the hydrophilic nature of the membrane, possibly due to the loss of the capsule (Fig 7C).

This was followed up using transmission electron microscopy to observe changes in the outer-membrane. *P. gingivalis* has an electron dense outer-membrane region that contains, among many components, the capsular polysaccharides. We observed that there were significant changes in the outer-membrane of the V3128 TonB deficient strain. The WT and complemented strains retained the electron dense region, however, the V3128 completely lacked this region (Fig. 7D).

## Discussion

Here we report on identification and characterization of a TonB energy transducer in *P. gingivalis*. Our data show that it is required for a variety of biological processes in *P. gingivalis*, including metal transport, black pigment accumulation, growth in broth media, protease activity and capsule formation. This is in agreement with the unprecedented lack of iron-dependent regulation of the TonB-coding gene^4 34^. Such a wide array of biological processes affected by the absence of TonB are evidenced by the extensive alteration of the protein profile of the TonB-deficient bacterial cell. The TonB energy transducer has been mainly characterized in Gram-negative aerobic bacteria and not much is known regarding the structure and role of the protein in anaerobic bacteria belonging to the Bacteroidota phylum^35 36^. This is a highly diverse group and while many of the gut Bacteroidota are saccharolytic and many utilize polysaccharides for their nutrition^37 38^, the *P. gingivalis* bacterium relies mainly on peptides derived from amino acids for its growth^9^. Indeed, the novel peptide transport system has been shown to be required for virulence of the bacterium^10 39 40^. Furthermore, the essential nutrient, iron, is also transported across the outer membrane in the form of heme and thus requires active transport. The above indicates the indispensable role of TonB in the biology of *P. gingivalis*. Given several unique properties of the *P. gingivalis* environment that includes being part of a biofilm and thus competition for nutrients with other bacteria in the biofilm as well as being an asaccharolytic bacterium and thus reliance on proteins for growth distinguishes it from bacteria such as *E. coli* where the TonB energy transducer was well characterized.

Our bioinformatics analyses show significant sequence similarity within the CTD domain. That also includes conservation of the critical functional amino acids. However, limited sequence similarity is present in the TM and LCCR domains indicating genus-specific characteristics of the protein. Some overlap was observed between the TM domains of *P. gingivalis* and *B. fragilis* TonB, however, sequence divergence was noted in the LCCR domain despite strong conservation of the genomic locus between the two bacterial genera. The conservation of the genomic locus composed of TonB and its accessory proteins ExbBD indicates that this locus may have a common ancestor.

Our mutant presented with a drastic phenotype. Given the essentiality of TonB such differences in phenotype are not surprising. It is indeed fortunate that we were able to identify conditions to generate a mutant deficient in TonB. The mutant grew well on blood agar plates which indicates that possibly a secondary TonB-like protein is operational under those conditions. Indeed, we identified the PG0633 as potential TonB protein. However, the PG0785 encoded TonB is essential for *P. gingivalis* growth in BHI broth. Its presence in *P. gingivalis* membrane has been shown by the Reynolds lab as complexed with the ExbB and ExbD proteins^41^ while the PG0633 has not been identified possibly pointing out to the importance of the complex for growth and fitness of the bacterium.

Our observations that the TonB-deficient mutant, *P. gingivalis* V3128 did not accumulate black pigmentation on the cell surface when grown on blood agar plates is in line with the observation of reduced gingipain activity in that strain. The Lys-specific gingipain has been demonstrated to be required for the pigment accumulation^7^ as the protease efficiently degrades hemoglobin and provides the bacterium with heme. Also, the Arg-gingipain has been shown to play a role in heme acquisition in *P. gingivalis* ^3^.

We also observed significant reduction in the ability of the mutant strain to survive in the presence of host cells. Such data are consistent with the lack of the µ-oxo-dimer forming the black pigmentation on the surface of the bacterium^3 42^ that would protect it from oxidative and nitrosative species generated by host cells in response to infection. Furthermore, the alteration in metal profile, especially reduced levels of manganese that plays a role in survival of the bacterium with host cells may have played a role ^43^. Ultimately, there is a large array of proteins that have differential expression in the mutant strain when compared with the parental one. It is evident that multiple factors led to reduction of fitness of the bacterium when exposed to host cells.

It is noteworthy that iron homeostasis was not affected but rather cell surface remodeling. The TonB was mainly reported to play a role in nutrient acquisition while the Tol system has versatile functions in cell outer membrane remodeling processes^44 45 46^. Such data indicates that there is possible functional overlap between the TonB and Tol motors^47 48^. Our functional analyses were guided by our proteomics data showing that the mutant strain has significantly downregulated levels of the capsule regulator tyrosine kinase locus (Ptk1)^1^. Indeed, we did find that the TonB-deficient mutant cells had lower levels of capsule that in turn could lead to reduced survival and elevated invasion of host cells^49^.

In conclusion, we were able to demonstrate that PG0785-encoded protein is important in binding of heme to the surface of *P. gingivalis* and formation of the black pigmentation when bacteria are grown on blood agar plates. We were also able to show that the PG0785-encoded TonB plays an important role in uptake of metals, protease activity and virulence of *P. gingivalis.* The *P. gingivalis* V3128 strain contains significantly less iron than parental W83 in bacteria grown on blood agar. This reduction of iron levels can be attributed to the abrogation of heme on the surface of the bacterial cells. The drastically reduced protease activity would also explain the reduced heme acquisition and virulence of *P. gingivalis*^3 7 50 51 52^. The proteomic analysis of *P. gingivalis* V3128 strain allows us to further assess the vast extent to which TonB contributes to the multitude of biological processes in *P. gingivalis* and thus deserves further in-depth investigation of its role in more *in vivo* settings as well as determination of the motor’s molecular structure and mechanism that would allow to design interventional strategies.

## Data Availability

The mass spectrometry proteomics data have been deposited to the ProteomeXchange Consortium via the PRIDE^53^ partner repository with the dataset identifier PXD033524.

**Project Name:** Role of TonB in alteration of protein profile of *Porphyromonas gingivalis*

**Project accession:** PXD033524

## Acknowledgements

This work was funded by NIH grants 1R01DE023304 (J.P.L.), NIHR21DE025555 (J.P.L.) and K18 DE029865 (J.P.L.). Services and products in support of the research project were generated by the VCU Massey Comprehensive Cancer Center (MCCC) Microscopy Shared Resource supported, in part, with funding from NIH-NCI Cancer Center Support Grant P30 CA016059. Figures were generated using the BioCyc Pathways Tools version 26.0 (software by SRI International).

We thank Qin Gui for her help preparing bacterial samples for proteomics analysis. The whole bacterial proteomics analysis was done by the John Hopkins University Center for Proteomics Discovery; we acknowledge the expert help from Ruiqiang Chen and Chan-Hyun Na. We also thank Dr. Tytus Bernas and Terry Smith of the VCU microscopy core for their help with design and analysis of the bacterial structures using microscopy approaches and electron microscopy. Finally, we thank Enila Hoxhalli for generation of constructs for PG0785 mutagenesis and Nicai Zollar for her help with the HL60 cell infection assays.

A portion of the work has been included in the MS thesis of the first author^54^.

## Author Contributions

S.R. performed experiments and wrote the manuscript. J.P.L. performed experiments, analyzed data, wrote and edited the manuscript. B.R.B. performed experiments, edited manuscript and figures.

## Supplemental Figures

**Figure S1. Blast search analysis of PG0785.** A. Sequence comparison of PG0785 with the *B. fragilis* 638R *tonB3*. B. List of the 11 most similar sequences to the PG0785.

**Figure S2. *P. gingivalis* TonB is required for sustained growth in broth cultures.** Wild Type W83, V3128 *tonB* mutant, and V3128 complement were grown for multiple passages in supplemented tryptic soy broth (TSB). Cultures were initiated from blood agar plates and 1:10 dilutions were performed for each passage. P1 – passage 1, P2 – passage 2, P3 – passage 3.

**Figure S3. *P. gingivalis* TonB is required for secreted gingipain activity.** Wild type W83, V3128 *tonB* mutant, and V3128 complement were grown for P1 in supplemented tryptic soy broth. Cultures were initiated from blood agar plates. The supernatant protease activity of Arg-gingipain (Panel A) and Lys-gingipain (Panel B) was chromogenically measured over 180 minutes using 1mM BApNA (RGP) or 0.5mM TGPLpNA (KGP). The x-axis represents the time in minutes of the measurement’s recordings. The y-axis represents the activity of proteases as assessed by measuring absorbance at 405 nm.

**Figure S4. *P. gingivalis* TonB affects metal content as determined using an ICP-MS analysis.** Bacteria (W83 and V3128) were grown on blood and BHI agar plates, bacterial cells were then suspended in 4 mls of BHI, pelleted by centrifugation at 7500 rpm for 10 minutes, washed twice with 4 mls of Chelex Buffer (0.05 M HEPES, 0.05 M NaCl), and suspended in 3 mls Chelex Buffer with 8 M Urea. Following incubation for 1 hour at room temperature bacterial cells were lysed using the sonication. The mixture was centrifuged at 14,800 rpm for 30 minutes and the supernatant was collected for metal content analysis. Samples were analyzed using an ICP-MS to determine the amount of iron (Fe), magnesium (Mg), manganese (Mn), zinc (Zn), nickel (Ni), copper (Cu) and cobalt (Co) in each sample. Metal content results for Ni, Cu, Zn and Co are shown in Panel A and results for Fe, Mg and Mn are shown in Panel B.

**Figure S5. TonB deficiency alters the protein profile of *P. gingivalis* cells.** A. V3128 (lane 1) and W83 (lane 2) cell lysate proteins derived from bacteria grown on blood agar plates resolved on SDS-polyacrylamide gel. Bands 1, 5, 6, 9, 12 and 14 from lane 1 and bands 1, 5, 6, 9, 19 and 14 from lane 2 were excised from the gel and sent for mass spectrometry analysis. B. Proteins identified from gel slices using Sequest search algorithm.

**Figure S6. Structure of genomic loci downregulated upon mutation of TonB in *P. gingivalis* W83.**

**Figure S7. Sample gating strategy.** Debris from cells and PBS and other small events were gated out. To ensure only cells were analyzed, the P1 gate was further gated for the DNA binding dye Syto17 to further delineate any debris from cells. This P2 gate was then measured for FITC fluorescence to assess the levels of polysaccharide capsule on the cell surface.

## Supplemental Tables

**Supplemental Table 1 S1.** All identified proteins in this study.

**Supplemental Table 2 S2.** V3128 vs W83 on blood agar plates.

**Supplemental Table 3 S3.** V3128 vs W83 on BHI plates.

**Supplemental Table 4 S4.** Statistically significantly regulated in W83 grown on blood agar vs BHI plates.

